# Bayesian model-based method for clustering transcriptomics time series with multiple replicates

**DOI:** 10.1101/2024.05.23.595463

**Authors:** Elio Nushi, François P. Douillard, Katja Selby, Miia Lindström, Antti Honkela

**Affiliations:** Department of Computer Science, Faculty of Science, University of Helsinki, Helsinki, Finland; Department of Food Hygiene and Environmental Health, Faculty of Veterinary Medicine, University of Helsinki, Helsinki, Finland

**Keywords:** Transcriptomics time series, RNA-Seq, TPM measure, Clustering, Gaussian process, Dirichlet process, Experimental replicates

## Abstract

**Background:** In this study, we introduce a Bayesian model-based method for clustering transcriptomics time series data with multiple replicates. This technique is based on sampling Gaussian processes (GPs) within an infinite mixture model from a Dirichlet process (DP). Our method uses multiple GP models to accommodate for multiple differently behaving experimental replicates within each cluster. We call it multiple models Dirichlet process Gaussian process (MMDPGP) mixture model. We compare our method with state-of-the-art model-based clustering approaches for handling gene expression time series with multiple replicates.

**Results:** To demonstrate the accuracy of our method we use it to cluster synthetically generated data sets. The clusters produced by our method on the synthetic data had a significantly higher purity score compared to the state-of-the-art approaches. We also present a case study where all methods are applied for clustering RNA-Seq time series of *Clostridium botulinum* with three different experimental replicates. The results obtained from the gene enrichment analysis showed that the number of significantly enriched sets of genes is larger in the clusters produced by MMDPGP.

**Conclusions:** By modelling each replicate with a separate GP, our method can use the natural variability between experimental replicates to learn more about the underlying biology.

## 1 Introduction

Clustering genes based on their transcription dynamics can yield invaluable information about the roles of these genes in different biological processes and their complex interaction network. The transcription dynamics are captured by gene expression time series, which can be collected in multiple replicates to take in consideration the biological variation and measurement errors [1].

Traditional clustering methods such as k-means, hierarchical clustering or self-organizing maps rely on correlation or Euclidean distance, which may not account for dependencies in gene expression levels across time points [2]. Alternatively, model-based clustering methods assume that the data are generated from a mixture of underlying probability distributions, each representing a different cluster [3]. However, previously published model-based clustering methods for gene expression time series have put little to no attention in considering the variability of experimental replicates. In this work, we present a Bayesian model-based clustering technique for multi-replicate transcriptomics time series, that uses a Dirichlet process (DP) [4] for generating clusters and multiple Gaussian process (GPs) [5] models to represent the different replicates inside each cluster. We call our method multiple models Dirich-let process Gaussian process (MMDPGP). This approach incorporates a measure of uncertainty to the cluster assignments. It also provides the added benefit of automatically identifying an optimal number of clusters based on different optimality criteria, instead of arbitrarily setting it to a fixed value like in the case of k-means clustering for instance.

We compare MMDPGP with two similar model-based approaches proposed by Hensman et al. [6] and McDowell et al. [2]. The approach presented by Hensman et al. is called mixture of hierarchical Gaussian processes (MOHGP) and it models the replicates using a single GP model at each cluster and samples the clusters from a DP. The software implementation of MOHGP indicated in the original Hensman et al. paper [6] was based on a variational inference technique. In order to provide a better comparison we implemented it using a Gibbs sampler written in Python similarly to [2]. On the other hand, McDowell et al. handle multiple experimental replicates by using the average of the replicates at each time point as the data in their GP model. They call their method Dirichlet process Gaussian process (DPGP). Unlike these alternative methods, MMDPGP provides a finer and more detailed modelling of each replicate which could help uncover clusters of genes that always behave similarly with each other, but together may behave differently across different replicates because of natural variability.

We use the methods for the purpose of clustering recently collected RNA-Seq time series. The RNA-Seq data were obtained by collecting RNA samples from growth experiments of three replicate cultures of *Clostridium botulinum* strain ATCC 19397 in different time points.

## 2 Related Methods

Time-series clustering has been extensively studied in statistics and data mining, with applications in many different domains. Several works provide general overviews of clustering methods specifically designed for time-series data, and discuss how these methods can be applied across fields sciences [7–9]. In this work, we focus on time-series clustering in the context of transcriptomics, where the aim is to group genes according to the similarity of their expression dynamics over time.

In the context of transcriptomics, clustering gene expression patterns according to their similarity has been the subject of many studies [2, 6, 10–12]. Rasmussen et al. [10] proposed a Dirichlet process mixture (DPM) model as an alternative to the bootstrap approach for modeling uncertainty in gene expression clustering. This method calculates the likelihood of two genes being in the same cluster as a measure of similarity. It does this by considering the primary factors of model uncertainty, such as the number of clusters and their location. They used a Gibbs sampler to implement their model and apply it to cluster high-dimensional non-time series gene expression data.

Medvedovic et al. [11] developed different clustering procedures based on Bayesian mixture models. Their method design specifically targeted microarray gene expression data with experimental replicates. They model the distribution of their data as a Bayesian mixture model and consider both finite and infinite Bayesian mixture models in their modeling approach. However, they do not present any application of their method on time series data.

Cooke et al. [12] introduced an extension to the Bayesian hierarchical clustering (BHC) algorithm [13] for use with time series data, and tested it on microarray time series. BHC uses a one-pass, bottom up greedy approach which assigns each data point to a separate cluster and iteratively merges cluster pairs together similarly to traditional agglomerative clustering. Their algorithm utilizes GP regression to model the structure of the data and can accommodate multiple replicate measurements of genes to enable a better discrimination of gene expression profiles.

Hensman et al. [6] introduced a clustering method which they called mixture of hierarchical Gaussian processes (MOHGP), that is based on GPs and DP. Their method can cluster time series with multiple replicates, but they consider only one way of modeling replicates which consists of clusters that are parameterized by a single GP. They provide an efficient implementation for MOHGP based on a variational inference procedure.

McDowell et al. [2] presented a method for clustering gene expression time series by combining a DP with a GP, and they called it simply DPGP. Their method is specialized in clustering gene expression time series consisting of a single replicate. If multiple replicate measurements are provided, it simply calculates their average and uses that in the rest of the analysis, thus not considering the variability between replicates.

In this work we focus on non-parametric clustering methods for time series which can handle the information provided by different replicates. Here we propose MMDPGP, a method for clustering gene expression time series which leverages the information provided by the variability between different experimental replicates. We compare our method with two previously published model-based clustering methods: MOHGP (Hensman et al.) and DPGP (McDowell et al.). Similarly to

MMDPGP, MOHGP considers the variability between experimental replicates. However, it assumes that the replicate gene measurements within the same cluster are generated by the same cluster-specific GP model, while MMDPGP assumes that the different gene expression replicates within the same cluster are generated by replicate-specific GPs. On the other hand, DPGP (McDowell et al.) ignores the variability between the replicates and uses only their average to fit the GPs in the clustering process. We argue that MMDPGP is superior to the previous methods for clustering gene expression time series with multiple replicates. We demonstrate this through different analyses after applying the methods on both biological data consisting of RNA-Seq time series of *C. botulinum* cultures and synthetically generated data. A short summary of the differences between our method and other clustering methods mentioned in this paper can be found in Table 1.

**Table 1:**
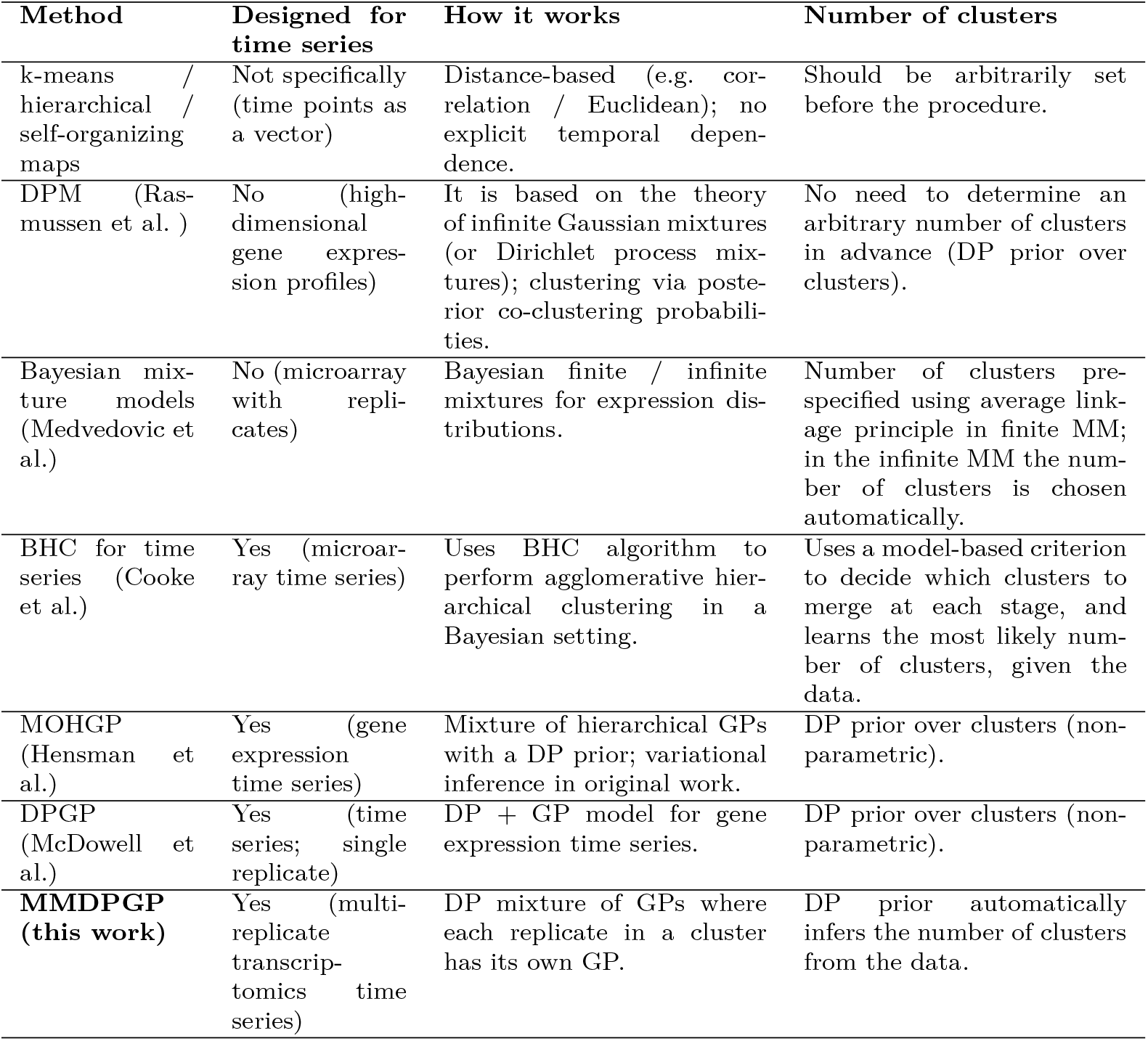
Comparison between different clustering methods.

| Method | Designed for time series | How it works | Number of clusters |
|---|---|---|---|
| k-means / hierarchical / self-organizing maps | Not specifically (time points as a vector) | Distance-based (e.g. cor-relation / Euclidean); no explicit temporal depen-dence. | Should be arbitrarily set before the procedure. |
| DPM (Ras-mussen et al. ) | No (high-dimensional gene expres-sion profiles) | It is based on the theory of infinite Gaussian mixtures (or Dirichlet process mix-tures); clustering via poste-rior co-clustering probabili-ties. | No need to determine an arbitrary number of clusters in advance (DP prior over clusters). |
| Bayesian mix-ture models (Medvedovic et al.) | No (microarray with repli-cates) | Bayesian finite / infinite mixtures for expression dis-tributions. | Number of clusters pre-specified using average link-age principle in finite MM; in the infinite MM the num-ber of clusters is chosen automatically. |
| BHC for time series (Cooke et al.) | Yes (microar-ray time series) | Uses BHC algorithm to perform agglomerative hier-archical clustering in a Bayesian setting. | Uses a model-based criterion to decide which clusters to merge at each stage, and learns the most likely num-ber of clusters, given the data. |
| MOHGP (Hensman et al.) | Yes (gene expression time series) | Mixture of hierarchical GPs with a DP prior; variational inference in original work. | DP prior over clusters (non-parametric). |
| DPGP (McDowell et al.) | Yes (time series; single replicate) | DP + GP model for gene expression time series. | DP prior over clusters (non-parametric). |
| **MMDPGP (this work)** | Yes (multi-replicate transcrip-tomics time series) | DP mixture of GPs where each replicate in a cluster has its own GP. | DP prior automatically infers the number of clusters from the data. |

## 3 Methods

### Gaussian Processes

Gaussian processes are non-parametric methods that are used for building probabilistic models of functional data [5]. In many previous studies, GPs have been widely recognized and adopted for modeling transcriptomics data [1, 2, 14–19]. A GP is completely characterized by a mean function *m*(*t*) and a covariance (or kernel) function *k*(*t, t*^*′*^), which is often referred to as the kernel of the GP. It defines a probability distribution over functions *f* (*t*) which we denote as:

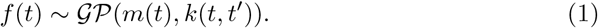

Choosing the kernel is an important step in the process of modeling with GPs as it determines our assumptions about the function that we want to model. The most widely used kernel function for modelling with GPs is the exponentiated quadratic function [5] which is defined as follows:

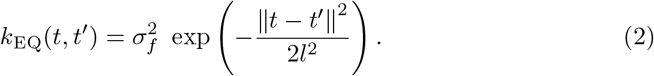

The exponentiated quadratic kernel function is characterized by 2 parameters which are the length scale *l* and the variance 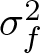. The length scale determines the oscillatory behavior of the function, specifically influencing the extent of fluctuations. Generally, the predictive capability of the model is limited to a range not exceeding *l* units from the observed data, indicating constraints on the model’s extrapolation potential beyond this threshold. The variance determines the average distance of the function away from the mean. We will also consider a third parameter 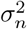 which is the Gaussian noise and represents the observation noise of the data in the GP model. In the rest of the analysis when we write *k*(*t, t*′) it is implied the exponentiated quadratic kernel unless specified otherwise. Our method uses GPs as a parametrization for each cluster of genes (i.e., all genes within the same cluster share the same GPs’ profiles).

### Infinite mixtures of Gaussian processes

Bayesian nonparametric (BNP) techniques differ from traditional methods in addressing the complexity of models. In standard cluster analysis methods like k-means, it is required to evaluate a range of models, each with a different parameter value (e.g., *k* = 1, …, *N* clusters), then the optimal model is selected based on various metrics for model comparison. Traditional methods are characterized by a finite number of parameters. In contrast, BNP methods are grounded in a statistical framework that can accommodate a potentially infinite number of parameters, this characteristic makes them flexible and removes the burden of choosing the number of parameters which could best model the system.

The Dirichlet process (DP) [4] is an example of a BNP technique which we use in our clustering method. DP is a nonparametric probability distribution over probability distributions, and it is used as a flexible prior for unsupervised learning tasks like clustering and density modeling [20]. We denote a probability distribution sampled from a DP as:

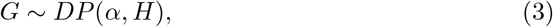

where *α* is a positive real number called the concentration parameter, and *H* is a probability distribution called the base measure.

In the context of clustering, a DP is used to generate priors for a Dirichlet process mixture model (DPMM) which is a mixture model that accounts for a theoretically infinite number of mixture components [21]. Several authors have extended the concept of DPMM to include GP models and call them infinite mixture of Gaussian process experts, infinite mixture of global Gaussian processes or Dirichlet process Gaussian process models [2, 6, 22–25].

In this paper we use a Dirichlet process Gaussian process model to cluster time course transcriptomics data using multiple replicates. Below we define the data and the model used for the clustering.

Let 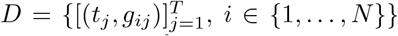, be a set of transcriptomics time series, where *g*_*ij*_ represents the gene expression value of gene *i* at time *t*_*j*_. We denote by *f*_*i*_(*t*) the latent function that models the time series 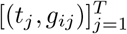 representing the gene expression values of gene *i* at every time point *t*_*j*_. Then, we define the following generative model for our data:

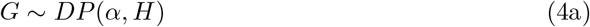

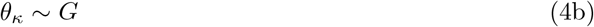

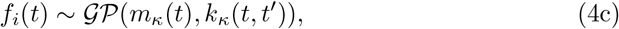

where *G* is a sample from *DP* (*α, H*) as defined in Eq (3) (small values of *α* generate fewer clusters with larger sizes, while large values generate more clusters with smaller sizes), and *θ*_*κ*_ = {*m*_*κ*_(*t*), *l*_*κ*_, *σ*_*f,κ*_, *σ*_*n,κ*_} is a set of GP latent parameters for cluster *κ*, which is drawn from the distribution *G*. Finally, we set *f*_*i*_(*t*) to be a sample drawn from the GP with a mean function 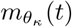, an exponentiated quadratic kernel parameterized by *l*_*κ*_, *σ*_*f,κ*_, and observation noise *σ*_*n,κ*_. It is possible to express the conditional distribution of a randomly sampled cluster parameter *θ*_*κ*_ dependent on all the previously sampled parameters *θ*_*\κ*_ by integrating over *G* and obtain:

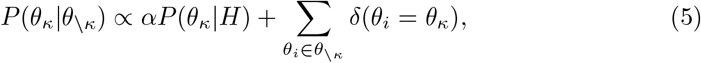

where *δ* is the Kronecker delta function, and *θ* _*κ*_ is the set of all sampled parameters except for *θ*_*κ*_. This suggests a practical solution to the problem of sampling from a DP because drawing a concrete distribution G is impossible in practice.

### Model selection for fitting multiple replicates

Gene expression time series can be collected in multiple replicates to take in consideration the biological variation and measurement errors [1]. However, the information provided by multiple replicates is often disregarded by clustering methods or they are used merely for estimating an average value which is later used in the downstream analysis as is the case in [2].

In this paper we construct and analyze 3 different approaches which take in consideration all replicate measurements of genes in the process of clustering genes by using a DP and GPs. The first method, MOHGP, was proposed by Hensman et al. [6]. The second method, DPGP, was proposed by McDowell et al. [2]. The third is our proposed approach, that we call multiple models DPGP (MMDPGP). Conceptually they involve changing Eq (4c) in the generative model defined by Eq (4a)–(4c). Below we explain the details for each approach.

To account for *R* different replicates, we extend the notation used for defining our transcriptomics time series data *D* in the previous section by introducing a superscript *r* (*r* ∈ {1, …, *R*}) which denotes the specific replicate time series (i.e., 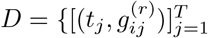, *i* ∈ {1, …, *N*}, and *r* ∈ {1, …, *R*}}).

### MOHGP (Hensman et al.)

In the case of MOHGP for each cluster *κ*, Hensman et al. [1] define a regression problem over functions *f*_*i*_ : *t* ↦ *f*_*i*_(*t*), which are generated by the following model:

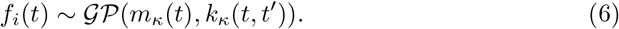

We denote by 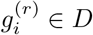 the vector 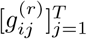 representing the gene expression values of gene *i* at every time point *t*_*j*_ of replicate *r*. The model assumes that they are noisy observations of *f*_*i*_(*t*). If we denote by 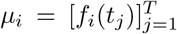 the vector representing the values of *f*_*i*_(*t*) at each time point *t*_*j*_, then the conditional distribution for 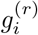 can be written as follows:

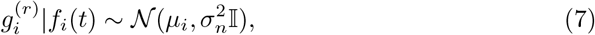

where 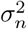 represents the noise variance between all replicates across all genes inside the cluster.

To make inference easier, *f*_*i*_(*t*) is marginalized out and the model is expressed by conditioning on *m*_*κ*_(*t*). In the marginalised model, the covariance between two observations 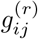 and 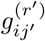 will then be

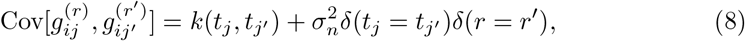

where *δ* is the Kronecker delta.

Let 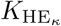 be the covariance matrix defined by the kernel function in Eq (8). If we denote by 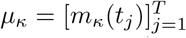, the vectors of observed values 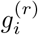 can then be sampled from the following multivariate normal distribution:

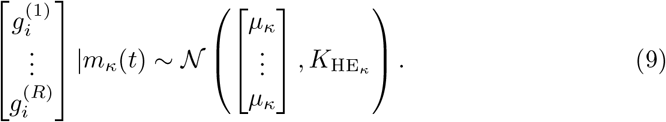

The value of *µ*_*κ*_ is determined as an approximation by setting it as 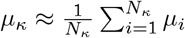, where *N*_*κ*_ is the number of genes in the cluster. The generative model MOHGP can be written as:

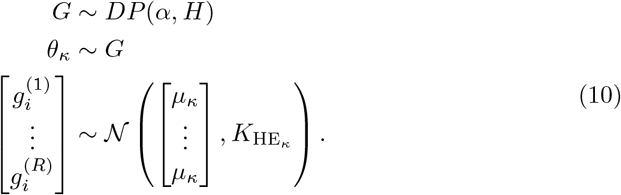

The computational complexity for fitting the GP of a single cluster in this model is therefore *O* (*N*_*κ*_(*RT*)^3^) in time and *O* (*N*_*κ*_(*RT*)^2^) in space, where *N*_*κ*_, *T, R* represent the number of genes in the cluster, the number of time points for which we have measurements and the number of replicates.

### DPGP (McDowell et al.)

Similarly to the previous approach, DPGP assumes that the latent mean function of each observed gene expression time series is sampled from cluster specific GP like in Eq 6. However, they limit their model by considering gene expression time series, which consist only of a single replicate. When multiple replicates are provided, DPGP collapses them into a single time series by computing their mean at each observed time point. Following the derivations of MOHGP from the previous section, but in the case of a single replicate, the generative model for DPGP can be written as:

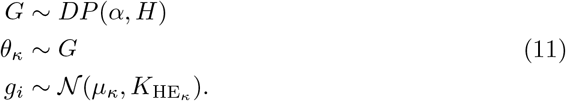

The computational complexity for fitting the GP of a single cluster for DPGP is *O*(*N*_*κ*_*T* ^3^) in time and *O*(*N*_*κ*_*T* ^2^) in space.

### MMDPGP

In the case of MMDPGP we take another approach by assuming that the set of functions *f*_*i*_ : *t* ↦ *f*_*i*_(*t*) are generated from replicate specific GPs within each cluster *κ* and we denote this association with superscript *r* 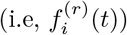. The generative model for 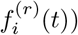 will then be

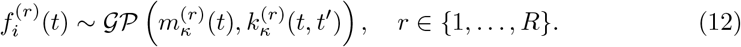

If we denote by 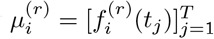 the vector of evaluations of 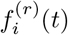 at time points *t*_*j*_, the vector 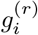 representing the gene expression values of gene *i* at every time point *t*_*j*_ of replicate *r* can be sampled from the following replicate specific conditional distribution:

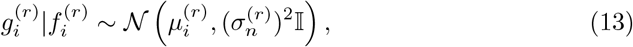

where 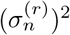 is the observation noise between genes of the same replicate in the cluster.

In order to perform inference on the model we can marginalize out 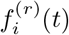 and express the model conditioned on 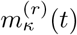. In the marginalized model the expression values of the same gene from different replicate measurements have a zero covariance, as they come from different GPs, we can reduce the computations by building smaller covariance matrices with the observations from each replicate separately.

In the MMDPGP model, each biological replicate *r* of a gene assigned to cluster *κ* is generated by its own latent function 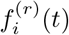, which is an independent draw from the replicate-specific GP of the same cluster. Thus, the expression values from different replicates are independent conditional to a common mean, and in the marginalized model, the prior covariance between observations belonging to different replicates of the same gene is zero, and the joint covariance matrix for 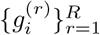 is block-diagonal with one block per replicate. This allows us to reduce computational cost by building and inverting smaller covariance matrices for each replicate independently. Importantly, replicates are still linked through their common cluster assignment and shared cluster-specific hyperparameters, so they are treated as multiple realizations of the same underlying temporal process rather than unrelated time series.

If we denote by 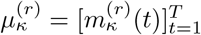, and by 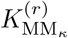 the covariance matrix in Eq (12) for genes in replicate *r*, we can sample 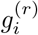 as:

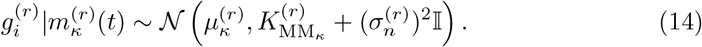

We can approximate the value of 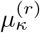 as 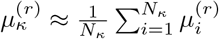 where *N*_*κ*_ is the number of genes in cluster *κ*. The generative model of MMDPGP will then be:

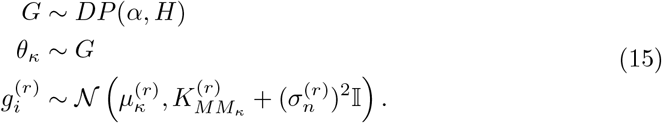

In typical transcriptomics experiments, one may encounter both biological and technical replicates. In our framework, biological replicates (e.g. independent cultures or individuals measured at the same time points) are modeled as distinct time series indexed by *r*, each with its own latent function 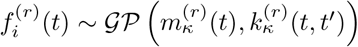 and kernel parameters 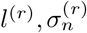. Variability between biological replicates is thus captured by the differences between these replicate-specific GPs within a cluster *κ*. In contrast, technical replicates (repeated measurements of the same sample) are not assigned separate GPs; they are modeled as additional noisy observations of the same latent trajectory via the replicate-specific observation noise 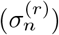 in Eq (13) and Eq (14).

The computational complexity for fitting the GP of a single cluster in the case of MMDPGP is *O*(*N*_*κ*_*RT* ^3^) in time and *O*(*N*_*κ*_*RT* ^2^) in space, which is significantly lower than MOHGP but larger than DPGP.

### Implementation

In this section, we focus exclusively on describing our implementations of MMDPGP and MOHGP (Hensman et al.). In the case of DPGP (McDowell et al.), we utilized their software, which is extensively detailed in [2].

In the Gaussian process Dirichlet process model (Eq (4a)–(4c)), the DP generates the distribution parameters 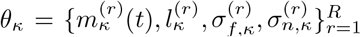 for the cluster specific GP(s). For 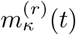, we denote by 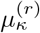 the vector of its evaluations on the observed time points. We set the following conjugate priors for the variance and observation noise as suggested in [2], which fit the characteristics of our data:

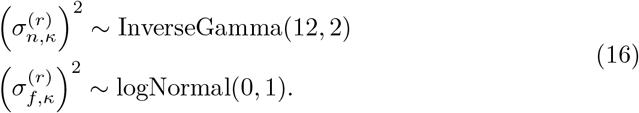

We set bounding constrains on the length scale 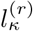 such that 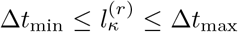, where Δ*t*_*min*_ = min(*t*_*i*_ *t*_*i* 1_) and Δ*t*_*max*_ = max(*t*_*i*_ *t*_*j*_) as described in [26], where *t*_*i*_, *t*_*j*_ are time points from the gene expression time series. Finally, we sample the means of the replicates 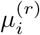 following Eq (6) and Eq (12) for MOHGP and MMDPGP accordingly. We initialise the parameters of each cluster with some default values (e.g., (*l*_0_, 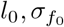, 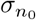) = (1, 1, 0.2)) to determine the initial covariance matrices 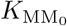, 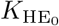 and for *µ*_0_ we sample a multivariate normal with zero mean, and covariance matrices 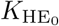 and 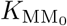 for each method respectively.

For the purpose of our clustering method we have used an implementation [2] of Neal’s Gibbs sampling Algorithm 8 [27], which yields the posterior probability for assigning a gene expression time series to a particular cluster. If we denote by *c*_*i*_ the cluster of gene *i* and *c\*_*i*_ the set of clusters of each gene except for gene *i*, the algorithm determines the probability for assigning gene *i* to any cluster *κ* conditioned on all other cluster assignments as follows:

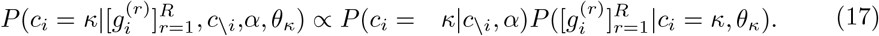

*P* (*c*_*i*_ = *κ c\*_*i*_, *α*) is a prior probability which does not depend on any characteristics of the time series for gene *i* but is based solely on the information about cluster *κ*:

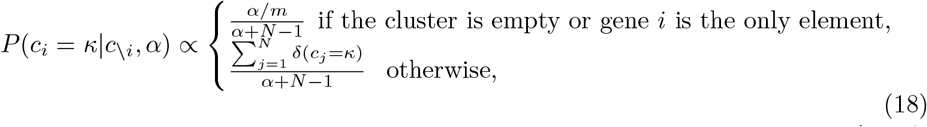

where *m* is the number of new clusters at each iteration which we set to 4, *α* is the clustering concentration parameter of the DP which we set to 1 as it has been found convenient on other gene clustering applications [28], *N* is the total number of clustered genes. However, while testing for different values of *α* and *m* we noticed that their impact on the number of clusters generated by MMDPGP is insignificant due to the good fit of the genes in the existing clusters. We provide more information about the procedure for choosing the optimal parameters *α* and *m* in section A.5 of the Appendix A.

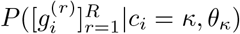 is the likelihood that the time series 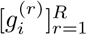 for all replicate measurements of gene *i* are generated by the GP(s) which parameterize cluster *κ*, and is given by Eq (19) and Eq (20) for MOHGP and MMDPGP respectively. Additionally, 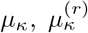 are calculated as averages, like we indicated in MOHGP and MMDPGP sections.

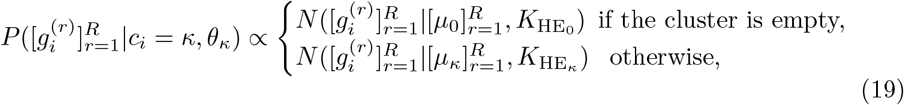

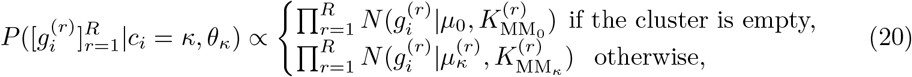

In order to select the optimal clusters for the genes, we use a posterior similarity matrix that indicates the proportion of samples generated by the Gibbs sampler for which every pair of genes (*i, j*) is found to be in the same cluster, as described in [2].

We assess the convergence of the clustering procedure using a trace plot for the number of occupied clusters (see section A.4 in Appendix A).

## 4 Data description and preprocessing

The data we used consisted of gene expression time series composed of RNA-Seq experiments sampled during different growth stages of *C. botulinum*.

### RNA-Seq data collection

The RNA-Seq data were obtained by collecting RNA samples from growth experiments of three replicate cultures of *C. botulinum* strain ATCC 19397 at 13 different time points. The three replicate cultures represent three biological replicates, as independent growth experiments performed under identical conditions. RNA samples were collected for each replicate at consistent time intervals corresponding to the growth experiment, specifically at 2, 3, 4, 5, 6, 7, 8, 9, 10, 11, 12, 15, and 24 hours post-initiation. Then, 92 ERCC RNA spike-in controls [29, 30] were added to the collected samples, and they were analyzed using paired-end sequencing with Ilumina’s NextSeq technology.

### Data preprocessing

Genes were assigned to COG (i.e., Clusters of Orthologous Groups of proteins) [31] categories using *EggNOG-mapper* [32] and were utilized in further analysis, as detailed below. We then carried out an operon identification analysis using *Rockhopper* [33].

*Operons* are clusters of co-expressed genes with related functions [34] and are often situated adjacently on the DNA strand, which leads to their collective activation. In the clustering analyzes presented here, we have used the information about operons as an informative prior by enforcing genes within the same operon to be in the same cluster. This approach would allow the GP(s) of each cluster to better capture the dynamic patterns of genes that are potentially involved in the same process, in what is called guilt by association [35].

During Gibbs sampling, we update the cluster label of each operon as a block: we sample a single cluster index for the operon and assign this index to all its member genes, with the likelihood given by the product of the gene-specific likelihoods under the candidate cluster. The same operon-based grouping is applied when running MMDPGP, MOHGP and DPGP, ensuring that all methods are compared on identical preprocessed data and constraints.

The gene expression data, which consist of values in TPM (i.e., transcripts per million) units, were then normalized using a spike-in-based normalization. Genes with low expression values were then filtered, as low values are not highly reliable and can introduce a potential source of noise in the analysis. A detailed description of the data preprocessing, including data preparation, normalization, and filtering, is included in sections A.1, A.2, and A.3 in Appendix A.

Before fitting the transcriptomics time courses to our model, we artificially pulled the last two time points of each time series closer to the rest—that is, the 15h and 24h time points were reassigned to 13.5h and 15h time points, respectively. This is because the model assumes that the transcription dynamics of the genes happened with the same intensity throughout the time course, which is not a realistic assumption in our case (i.e., in the beginning of the life-cycle [2h-12h] we expect rapid non-monotonic changes, while at the end the changes are very small and monotonic). Alternatively, the problem could be addressed by using a non-stationary kernel similarly to [36], but given that the change of transcription dynamic is expected only on the later time points we implemented this simpler approach.

## 5 Results

### Clustering of the genes using RNA-Seq time series

We applied the multi-replicate clustering methods MMDPGP, MOHGP [6] and DPGP [2] on the RNA-Seq data to group the genes according to their transcription dynamics. Even though we used the same values for initializing the parameters of the methods we obtained very different results. In the case of MMDPGP we acquired 26 different clusters, MOHGP method returned 116 clusters where most of the clusters have a small number of genes, while DPGP yielded only 13 clusters. The reason for this is because in the case of MOHGP during each iteration of the Gibbs sampler the likelihood of each gene to belong to a new cluster is higher. This is because there is only one GP for each cluster which captures the variation between all replicates, and given a not negligible difference between replicates, it often results in a poor fit for the gene under consideration, thus the gene is assigned to a new cluster. In the case of MMDPGP however, every cluster has a separate GP for each replicate and the biological replicates are compared on how well they fit in the replicate specific GP at each cluster, thus resulting in a better fit at the cluster whose genes express similarly on the same replicate. DPGP on the other hand ignores the variation between replicates and performs clustering using the replicates’ means. When performing the mean of the replicates, the replicate’s gene specific nuances vanish resulting in a larger number of similar time series, hence the smaller number of clusters. Fig 1 shows the GP models for three clusters generated by the methods. We can see how in the case of MMDPGP the GPs of different replicates differ because even though the replicates are similar they have their own specific characteristics depending on natural variation between the different replicates. When exploited, this natural variation can be a useful source of additional knowledge during the clustering procedure.

**Fig. 1:**
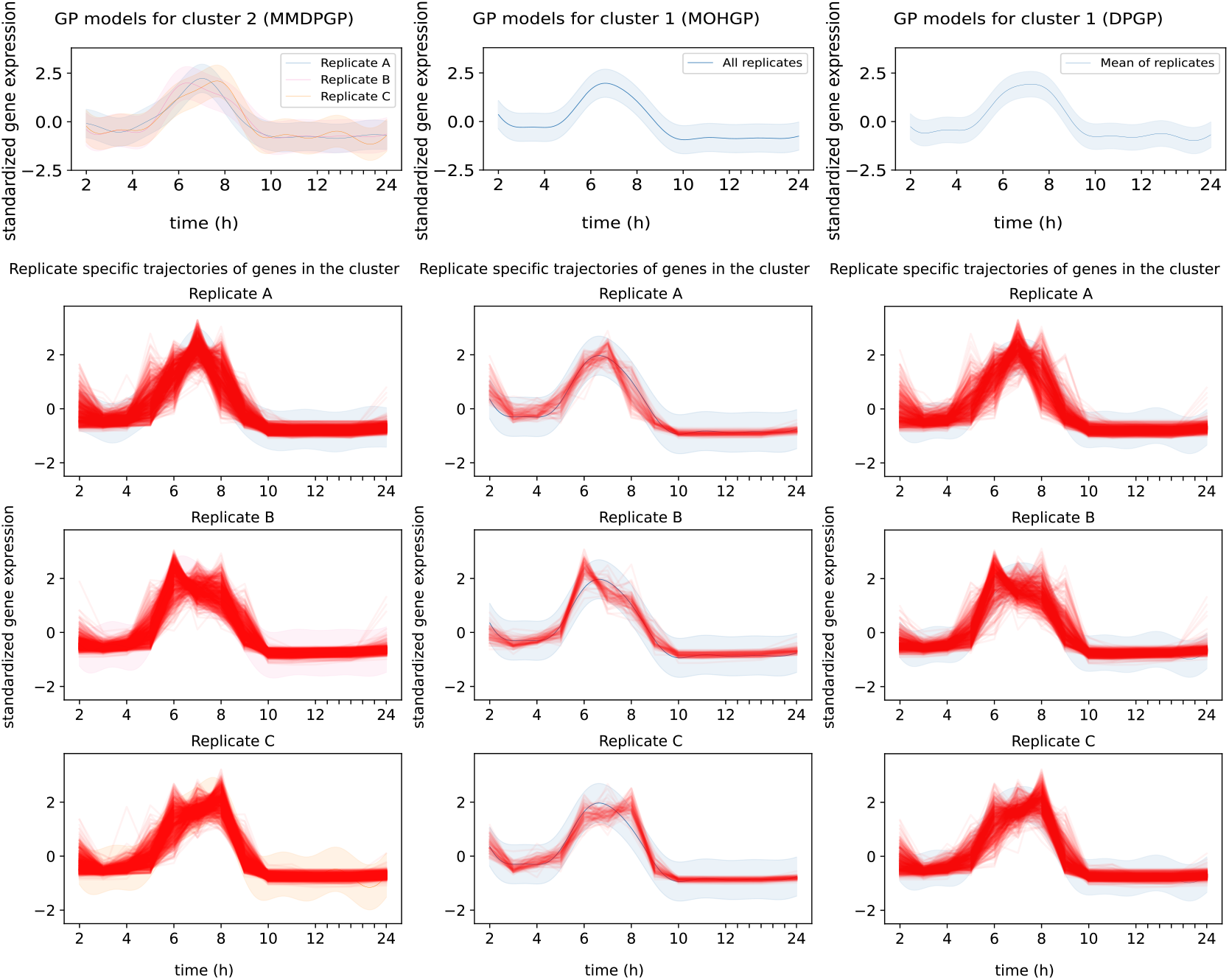
Comparison between 3 clusters generated by MMDPGP (left column plots), MOHGP [6] (middle column plots), and DPGP [2] (right column plots). The three replicates in the MOHGP’s cluster and DPGP’s cluster are modeled by one GP (top middle column plot and top right column plot). The three replicates in the MMDPGP cluster are modeled by three different GPs (top left plot). The replicate-specific trajectories of the genes in each cluster are plotted separately as red lines on top of their respective GP. In the case of MOHGP and DPGP they are plotted on top of the same GP model which characterizes the respective cluster. In the case of of MMDPGP cluster the trajectories are plotted on top of their respective replicate specific GP. The x-axis is drawn to indicate the real time of the RNA-Seq measurements.

### Biological significance analysis

To understand the biological relevance of our results, we carried out a gene enrichment analysis using COG annotations assigned to each gene (see “Data preprocessing” subsection). We chose this method because the standard gene ontology (GO) enrichment analysis, was not available for *C. botulinum* that is not annotated in the GO database. By using COG annotations, it is still possible to group the genes in a meaningful way and explore their potential functions.

To assess whether a given gene cluster is associated with specific functional categories, we performed an over-representation analysis using COG functional classes. For each cluster and each COG class, we built a 2×2 contingency table comparing the number of genes in the cluster that are annotated to that class with the number of genes outside the cluster that carry the same annotation. We then tested for enrichment using a hypergeometric test, and controlled for multiple testing across COG classes using the FDR procedure. In this formulation, the expected number of genes from a particular COG class in a cluster is determined by both the cluster size and the total number of genes annotated to that class in the full data set, so differences in COG class size are explicitly taken into account. COG categories are not strictly independent, and some classes may be related in a hierarchical manner. Our analysis treats each functional class as a separate gene set and does not attempt to model the dependency structure between classes explicitly. Thus, the resulting FDR-adjusted p-values should be interpreted as exploratory indicators of over-representation.

Fig 2 shows the significant COG enrichment across clusters obtained from each of the methods. The significant COG enrichment was determined by considering those which have a FDR-corrected *p*-value smaller than 0.05. The significantly enriched COG classes at each cluster are marked with −log_10_(*p*) while the rest are marked with 0. The level of significance for the significantly enriched COG classes is indicated by the number of stars in the heatmap.

**Fig. 2:**
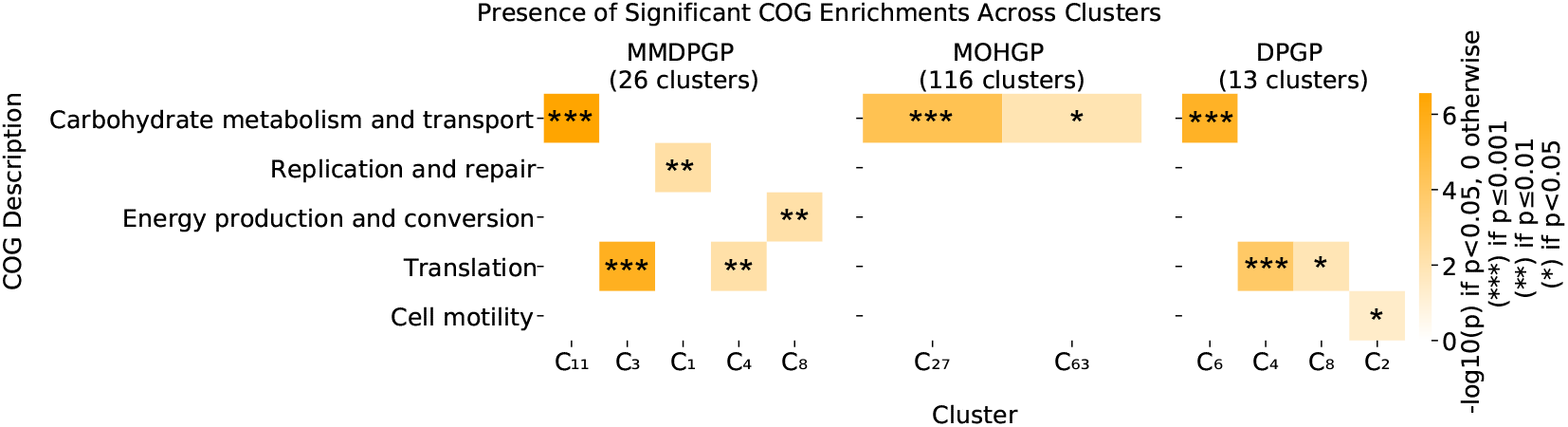
Gene enrichment analysis based on COG for MMDPGP (proposed), MOHGP [6] and DPGP [2]. Presence of significant COG enrichment is shown by −log_10_(*p*) for the COG categories with *p <* 0.05 at each cluster. COG categories which do not pass the *p*-value threshold are marked with 0. The enrichment significance of the COG classes is indicated by the number of stars in the corresponding class.

The heatmap for MMDPGP reveals a total of 5 significantly enriched COG classes and significant enrichment was detected in 5 clusters. In contrast, the heatmaps for MOHGP and DPGP methods show 2 and 4 detections of significant enrichments respectively. For higher levels of significance (i.e., *p* ≤ 0.01), MMDPGP still reveals 5 significantly enriched COG classes, while the numbers drop to 1 and 2 significant enrichments for the other methods. Additionally, all the significantly enriched COG classes with *p* ≤ 0.01, which are detected in MOHGP and DPGP, are also detected in MMDPGP. These indicate that the genes within each cluster of MMDPGP display patterns of higher biological similarity, compared to those inside the clusters that were generated by the other methods.

### Leave-one-out predictive likelihood

To estimate the quality of the fit of the genes in their corresponding cluster, we performed a leave-one-out predictive likelihood analysis as described in [37]. This is a standard measure for performing model selection with GPs. In order to do it, we refit the data of each cluster to its respective GP model by removing one time point from each gene/operon at a time. At each iteration, we calculated the log-likelihood of the model to predict the observed gene expression values at the time point that was removed. We did this for all time points (i.e., 13 time points) and we divided the numbers with the total number of single observations, to express the results in natural logarithm units per observation (nat/obs.). This normalisation allows for a fair comparison, given that in the case of DPGP the number of observations is 3 times smaller because they use only the mean of the 3 replicates as observation data. Fig 3 shows the results of the leave-one-out predictive likelihood analysis. We can see that MOHGP performs slightly better in the analysis by having a higher predictive likelihood for the time points overall compared to MMDPGP. This result can be explained by the fact that in the case of MOHGP most of the clusters have a very small number of genes which results in a better modeling of the data, but at the expense of producing less biologically meaningful clusters, as indicated by the gene enrichment analysis.

**Fig. 3:**
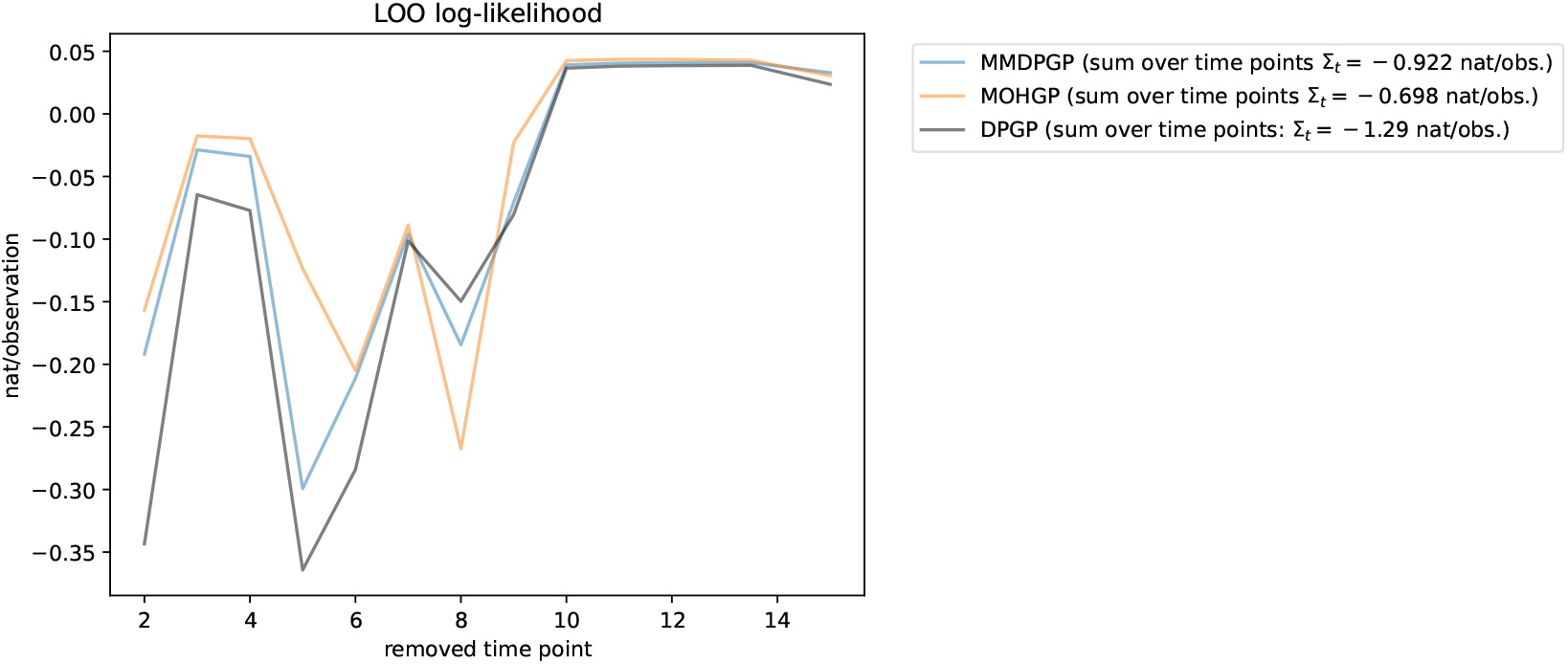
Sum of predictive log-likelihoods at each time point for the different methods (i.e., MMDPGP (proposed), MOHGP [6] and DPGP [2]) on the left out time points for each observation. Each point represents the sum of log-likelihoods of the observed gene expression values at the removed time point over all the posterior cluster models divided by the number of observations. In the legend we can also see the sums over all the removed time points, for each model.

### Performance on synthetic data with different amount of noise

We tested the classification accuracy of our methods by generating time series data from 6 different clusters. Each cluster was represented by a GP with a different mean function, and same kernel parameters (*l, σ*_*f*_) and observation noise 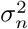. We sampled 10 different time series from each GP with mean *m*(*t*) ∈ {sin(*x/*2), −sin(*x/*2), sin(*x/*4), −sin(*x/*4), cos(*x/*2), −cos(*x/*2)} and introduced 2 additional replicates. For the first 5 clusters the replicates were generated by adding random noise *σ*. The additional replicates from the last cluster were obtained by horizontally shifting to the right (e.g., suggesting genes which are expressed earlier or later in different replicates) the initial samples by Δ and 2Δ respectively.

We simulated the process multiple times for different values of the GP model variance 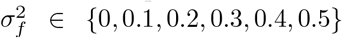, replicate noise standard deviation *σ* ∈ {0, 0.2, 0.4, 0.5}, and length of phase shift Δ ∈ {0, 0.3, 0.5}. The length scale and model noise were set to default values (i.e., 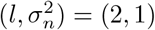). Each simulation resulted in clusters’ data with different attributes that are determined by the triple 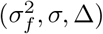. Then, we run the clustering methods with each cohort of generated data. In order to evaluate the results obtained by the methods we used a measure of purity for the clusters as defined in [38]. Purity assesses clustering quality by evaluating how well it avoids mixing different types of objects within the same cluster. This is a desired property in our context if we want to generate hypothesis about the functionality of unknown genes based on genes which we know in the same cluster. Fig 4 shows the purity values of the clusters obtained by running MMDPGP, MOHGP and DPGP methods on different cohorts of synthetic data. As it can be seen MMDPGP outperforms the other methods at all cohorts of synthetic data in terms of cluster purity measure. This means that MMDPGP is less likely to put time series which are originally generated by different clusters into the same one.

**Fig. 4:**
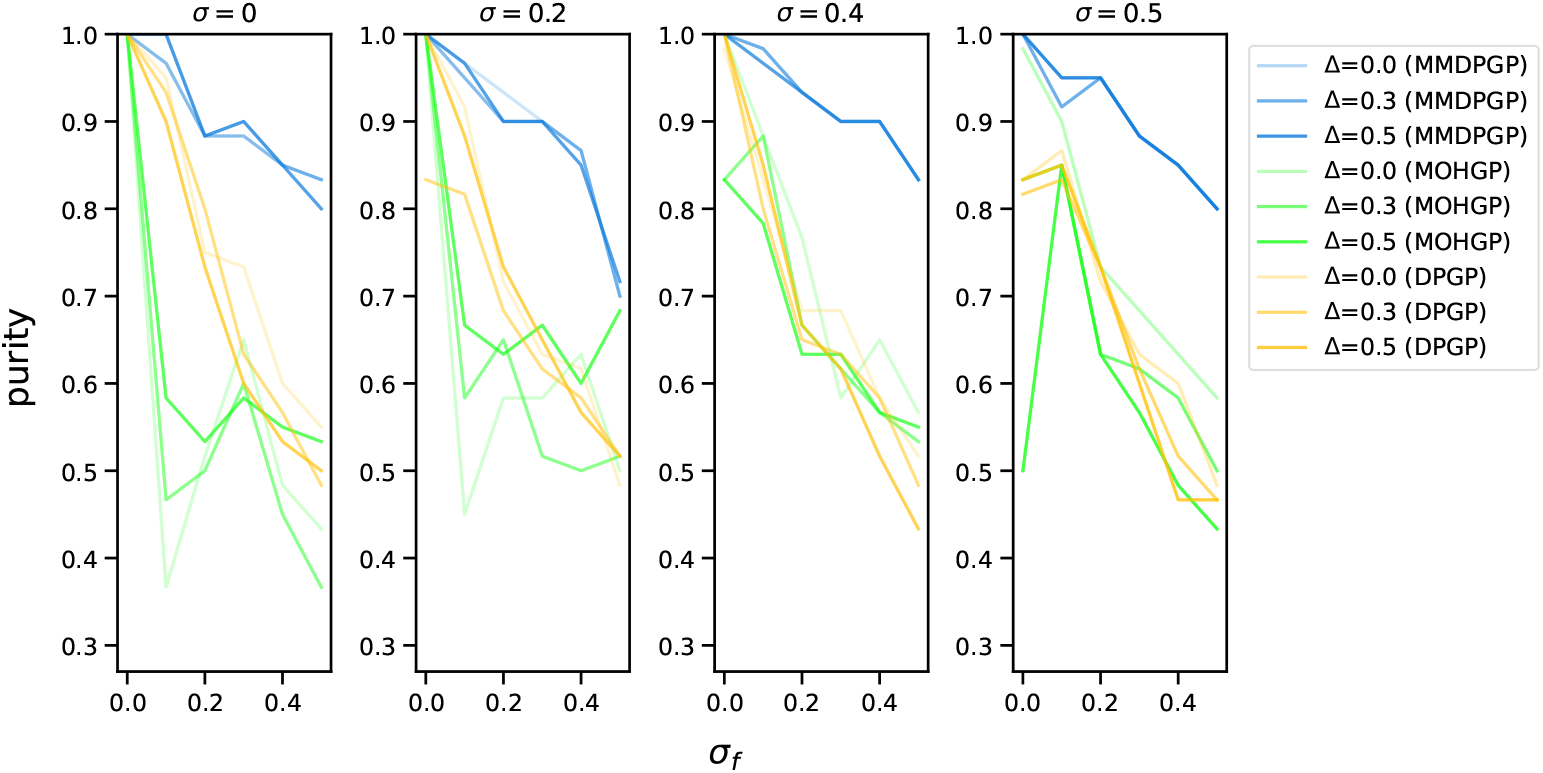
Cluster purity of the results obtained by applying MMDPGP (blue), MOHGP [6] (green) and DPGP [2] (amber) on synthetic data generated with different parameters (*σ*_*f*_, *σ*, Δ).

## 6 Discussion

Dirichlet processes (DP) and Gaussian processes (GP) can be combined and used to cluster functional data without making strong assumptions such as the number of clusters to group the data. In this paper we proposed a model-based method for the purpose of clustering gene expression time series with multiple replicates. Our method uses a different GP to represent each replicate inside every cluster, for this reason we call it multiple models Dirichlet process Gaussian process (MMDPGP) mixture model. We compared our method with two similar approaches, MOHGP, proposed by Henseman et al. [6], and DPGP, proposed by McDowell et al. [2]. We described the three methods and their generative model assumptions in detail. Python implementations were also provided for MMDPGP and MOHGP. One clear advantage of MMDPGP over MOHGP is its lower computational complexity which makes it useful for handling larger amount of data (e.g., data consisting of dense time series of thousands of genes, with many replicates). On the other hand even though DPGP has an even lower computational complexity compared to MMDPGP, it ignores the variation between replicates, which can be a useful source of information for clustering the genes in a biologically meaningful way.

We applied the clustering methods on a novel RNA-Seq time series data set consisting of three experimental replicates. We performed a COG enrichment analysis on the clusters produced by each method and showed that in the case of MMDPGP a higher number of significant enrichments (i.e., *p <* 0.05) are detected. Additionally for higher levels of significance (i.e., *p <* 0.01) not only the number of significant enriched classes is higher in the case of MMDPGP but all the significant enrichments detected by the other methods were also detected by MMDPGP.

The leave-one-out predictive likelihood analysis showed that MOHGP performed better than MMDPGP, but that is because the size of the clusters returned by MOHGP are very small which can cause an overfit to the GP of the clusters, thus yielding higher predictive likelihood overall.

Finally, to demonstrate the accuracy of MMDPGP we generated synthetic data from six different clusters by introducing different levels of noise and tested the performance of each method by measuring the cluster purity of the results. MMDPGP outperformed the other methods by scoring a significantly higher cluster purity as shown in Fig 4. We believe that MMDPGP can be useful for clustering gene expression time series data with multiple replicates because it effectively captures the information provided by experimental replicates and uses it to provide a meaningful discrimination between gene expression profiles. By modeling each replicate with a separate GP it provides more details for each individual replicate which can be useful for clustering in the cases when replicates differ, for instance because of different experimental conditions.

## Declarations

### 6.1 Ethics approval and consent to participate

Not applicable

### 6.2 Consent for publication

Not applicable

### 6.3 Availability of data and materials

#### 6.3.1 Data availability

The RNA-Seq data used to validate our method in the paper are deposited in GEO at the following link: https://www.ncbi.nlm.nih.gov/geo/query/acc.cgi?acc=GSE248529. The reviewers can access the data by providing the access token: *qfajmuisbvcxran*.

#### 6.3.2 Code availability

The Python code for implementing our method is publicly available and can be found on https://github.com/PROBIC/MMDPGP-clustering

### 6.4 Competing interests

We declare no competing interests related to this work.

### 6.5 Funding

The present study was supported by the European Research Council (ERC) under the European Union’s Horizon 2020 research and innovation programme (ERC-CoG whyBOTher, grant 683099), the HiLife Fellows Program and by the University of Helsinki. Open access funded by Helsinki University Library.

### 6.6 Author’s contributions

E.N. and A.H. developed the method; E.N. implemented the model and performed computational analyses; F.D., K.S., and M.L. provided the RNA-Seq data and biological expertise; E.N wrote the manuscript; all authors reviewed the manuscript.

## 6.7 Acknowledgments

We thank the Biomedicum Functional Genomics Unit at the Helsinki Institute of Life Science and Biocenter Finland at the University of Helsinki for performing the sequencing of the samples analyzed in this work. We are grateful to Hanna Korpunen for her excellent technical work in the laboratory.

## Notes

https://www.ncbi.nlm.nih.gov/geo/query/acc.cgi?acc=GSE248529

### Competing Interest Statement

The authors have declared no competing interest.

### Summary of Updates

This version of the manuscript has been revised to address reviewers' comments.

