## Appendix A for "Bayesian model-based method for clustering transcriptomics time series with multiple replicates"

### A.1 Data preparation

The paired-end raw sequencing reads generated from the sequencing phase were processed using RNA-Seq analysis software package *ProkSeq* [1]. ProkSeq performs quality control, filtering, and mapping to the reference genome of the reads. Then, it calculates the read counts for each gene or RNA spike-in and returns their expression values in TPM (i.e., transcripts per million) and CPM (i.e., counts per million) units. For the rest of the analysis we considered only the TPM gene expression values of the 13 different time points. This includes 3491 genes of *C. botulinum* ATCC 19397 genome and 92 ERCC RNA spike-in controls.

### A.2 Data normalisation for TPM gene expression time series

Zhao et al. [2] describes how TPM values are not always reliable to allow for cross-sample comparison of gene expressions. This is because TPM values represent the relative abundance of transcripts of each gene considering the full set of transcripts within the same sample [2]. Thus, comparing TPM values from different samples is done under the assumption that the total RNA concentration and distributions are very similar across samples [2]. This assumption is not always fulfilled as it is the case with our RNA-Seq samples. Fig A1 shows an artifact of the transcription dynamics of *BotR* (botulinum neurotoxin transcription-activating sigma factor) which is known to decrease at the late stages of the life cycle of the bacteria, but the TPM value increases instead. Therefore, to make our analysis more robust we conduct the following normalization method of the TPM values for each RNA-Seq sample using the ERCC RNA spike-in controls as described below.

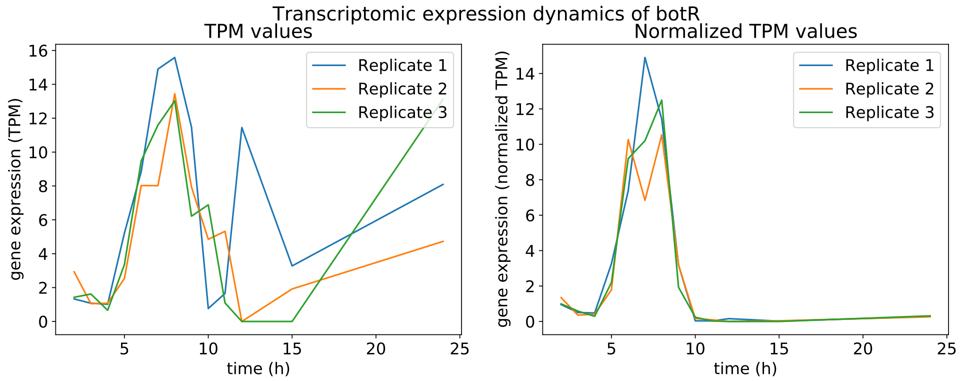


**Fig. A1**: Transcription dynamics of *botR* according to TPM values (left) and normalized TPM values (right).

Suppose our data consist of $N$ genes, $M$ spike-ins, and $K$ samples as depicted in Table A1. We denote by $g_{ij}$ and $s_{ij}$ the TPM value of the $i$th gene (spike-in) on the $j$th sample.

**Table A1**: Representation of the RNA-Seq gene expression data after quality control, filtering, and mapping to the reference genome of the reads (values are in TPM units)

| Gene/Spike-in | Sample$_1$ | Sample$_2$ | ... | Sample$_K$ |
|---|---|---|---|---|
| gene$_1$ | $g_{11}$ | $g_{12}$ | ... | $g_{1K}$ |
| gene$_2$ | $g_{21}$ | $g_{22}$ | ... | $g_{2K}$ |
| ... | ... | ... | ... | ... |
| gene$_N$ | $g_{N1}$ | $g_{N2}$ | ... | $g_{NK}$ |
| spike-in$_1$ | $s_{11}$ | $s_{12}$ | ... | $s_{1K}$ |
| spike-in$_2$ | $s_{21}$ | $s_{22}$ | ... | $s_{2K}$ |
| ... | ... | ... | ... | ... |
| spike-in$_M$ | $s_{M1}$ | $s_{M2}$ | ... | $s_{MK}$ |
| Sum | $10^6$ | $10^6$ | ... | $10^6$ |

Additionally, we have scaling factors $S_j$ which represent amount of spike-ins added on the $j$th sample as shown in Table A2. Given that spike-ins are added in varying

**Table A2**: Scaling factors showing the relative amounts of spike-ins added to each RNA-Seq sample.

| | Sample$_1$ | Sample$_2$ | ... | Sample$_K$ |
|---|---|---|---|---|
| Scaling factor | $S_1$ | $S_2$ | ... | $S_K$ |

quantities across samples, we need to normalize the data to ensure comparability between samples. To achieve this, the spike-ins should be adjusted by dividing them with appropriate scaling factors, thereby standardizing their levels and facilitating accurate comparisons.

First we partition the data into the set of genes and spike-ins and find $\mu = [\mu_1, \dots, \mu_K]$ such that $\mu_j \sum_{i=1}^{N} g_{ij} = 10^6$ for $j \in \{1, \dots, K\}$. We then transform the spike-ins $s_{ij}$ into $s'_{ij} = \mu_j \frac{s_{ij}}{S_j}$.

Given the assumption that the spike-ins are on the same amount in all the experiments, we fix the values of one column (say column corresponding to Sample$_1$) and for each column $j$ ($\forall j \in \{1, \dots, K\}$) we find a coefficient $c_j$ to multiply all the values of that column such that $c_j \cdot s'_{ij} = s'_{i1}$ ($i \in \{1, \dots, M\}$). In practice such a coefficient does not exist but we can choose it such that it minimizes the sum of squares of the differences:

$$\min((s'_{11} - c_j \cdot s'_{1j})^2 + (s'_{21} - c_j \cdot s'_{2j})^2 + \dots + (s'_{M1} - c_j \cdot s'_{Mj})^2),$$

and by solving it we find:

$$c_j = \frac{\sum_{i=1}^{M} s'_{i1} \cdot s'_{ij}}{\sum_{i=1}^{M} s'^2_{ij}}.$$

We multiply the gene expression data with the $c_j$ coefficients and obtain our normalised data as shown in Table A3.

**Table A3**: Normalised gene expression data using spike-ins. Notice how the TPM values of one column (in this case $\text{Sample}_1$) are left unchanged since $c_1 = 1$ and all the others are scaled relative to that.

| Gene/Spike-in | $\text{Sample}_1$ | $\text{Sample}_2$ | ... | $\text{Sample}_K$ |
|---|---|---|---|---|
| $\text{gene}_1$ | $c_1\mu_1 g_{11}$ | $c_2\mu_2 g_{12}$ | ... | $c_N\mu_N g_{1K}$ |
| $\text{gene}_2$ | $c_1\mu_1 g_{21}$ | $c_2\mu_2 g_{22}$ | ... | $c_N\mu_N g_{2K}$ |
| ... | ... | ... | ... | ... |
| $\text{gene}_N$ | $c_1\mu_1 g_{N1}$ | $c_2\mu_2 g_{N2}$ | ... | $c_N\mu_N g_{NK}$ |
| Sum | $10^6$ | $\neq 10^6$ | ... | $\neq 10^6$ |

In cases where the spike-ins are not reliable for a particular sample, we could deduce the coefficient $c_j$ by linearly interpolating in the log space from the coefficients of the samples belonging to the neighboring time points and set $c_j = \sqrt{c_{j-1}c_{j+1}}$.

### A.3 Data filtering

In the final step of the data preprocessing, we excluded all genes with a maximum expression level below 10 TPM across all time points. This decision was based on the fact that genes with low expression levels tend to have a minimal signal-to-noise ratio [3]. Consequently, these genes are more likely to add noise rather than meaningful information to our analysis.

### A.4 Convergence testing

In Fig A2 we show the trace plot of the number of clusters during each iteration of the sampler.

### A.5 Sensitivity analysis

We performed a grid search on different values of $\alpha = [1, 4, 6, 8]$ and $m = [1, 4, 6, 8]$. We ran the analysis on the full dataset composed of 2828 genes and 13 time points, and noticed that the number of clusters after the burn-in phase is very similar during the iterations of the sampler. We put the trace plots of the number of clusters for each iteration for all combinations $(\alpha, m)$ in Fig A3. For all pairs of $(\alpha, m)$, we select the optimal clustering based on the posterior similarity matrix and perform a leave-one-out log-likelihood analysis [4] to select the optimal pair $(\alpha, m)$ that maximizes the LOO log-likelihood. Fig A4 shows the LOO log-likelihood for the optimal number of clusters (according to the posterior similarity matrix) for different values of $(\alpha, m)$.

### A.6 Clustering the data after removing the last two time points

We run our clustering method using the same data after removing the last two time points. The first thing to notice is that the resulting number of clusters is 24 as

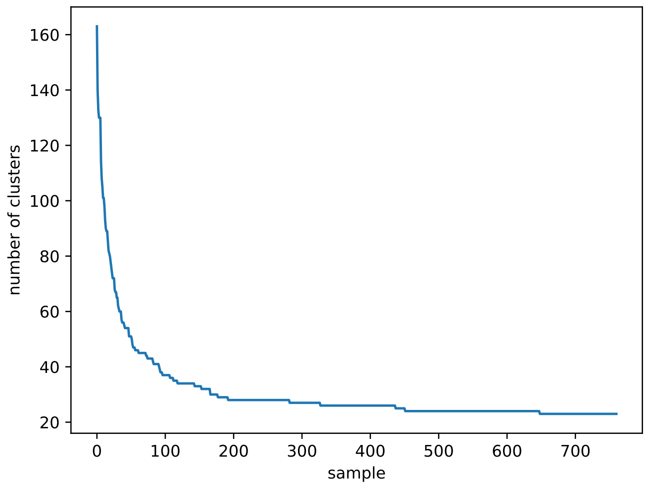


**Fig. A2**: Trace plot showing the number of occupied clusters during each iteration of the sampler after the burn-in phase.

**Table A4**: Comparison summary between the two clusterings.

| **Section** | **Quantity** | **Count** |
|---|---|---|
| Input summary | Aligned genes ($N$) | 2,828 |
| | # clusters in A (original dataset with all time points) | 26 |
| | # clusters in B (dataset without the last 2 time points) | 24 |

| **Pair counts** ($i < j$) | **Count** |
|---|---|
| Total pairs $\binom{N}{2}$ | 3,997,378 |
| $a$ same in both (A and B) | 365,371 |
| $b$ same in A only | 432,157 |
| $c$ same in B only | 184,542 |
| $d$ different in both | 3,015,308 |

| **Matrix metrics** | **Value** |
|---|---|
| Disagreement rate $\frac{b+c}{\binom{N}{2}}$ | 0.154276 |
| Rand index $\frac{a+d}{\binom{N}{2}}$ | 0.845724 |

opposed to 26 clusters that we obtained originally when using all the time points. In order to compare the similarity between the different clusterings we calculated the Rand index [5] which turned out to be approximately 85%, and can be interpreted as a measurement of agreement between the two different clusterings. Table A4 shows the summary of the results.

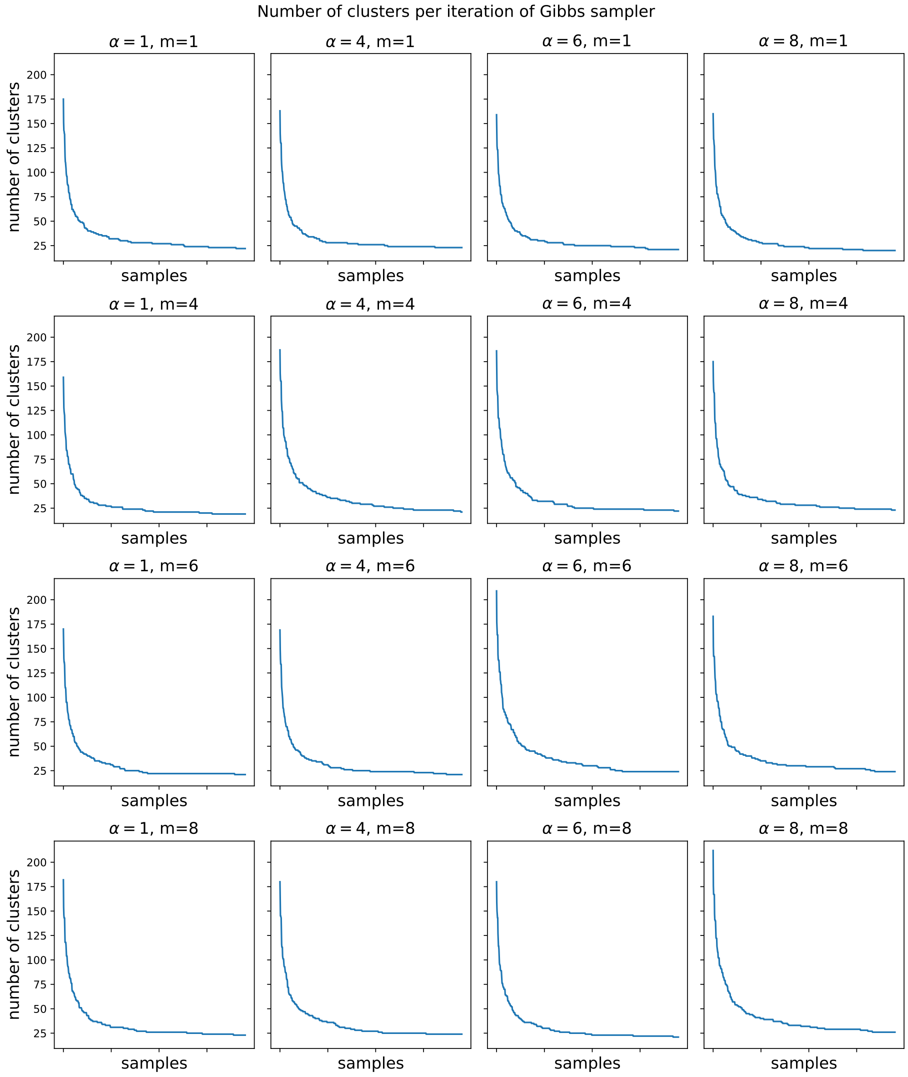


**Fig. A3**: Number of clusters per iteration of Gibbs sampler for different values of $(\alpha, m)$.

https://doi.org/10.1093/bioinformatics/btaa1063 https://arxiv.org/abs/https://academic.oup.com/bioinformatics/article-pdf/37/1/126/41921780/btaa1063.pdf

[2] Zhao, S., Ye, Z., Stanton, R.: Misuse of RPKM or TPM normalization when comparing across samples and sequencing protocols. RNA **26**(8), 903–909 (2020)

[3] Ng, V., Lin, W.-J.: Comparison of assembled Clostridium botulinum A1 genomes

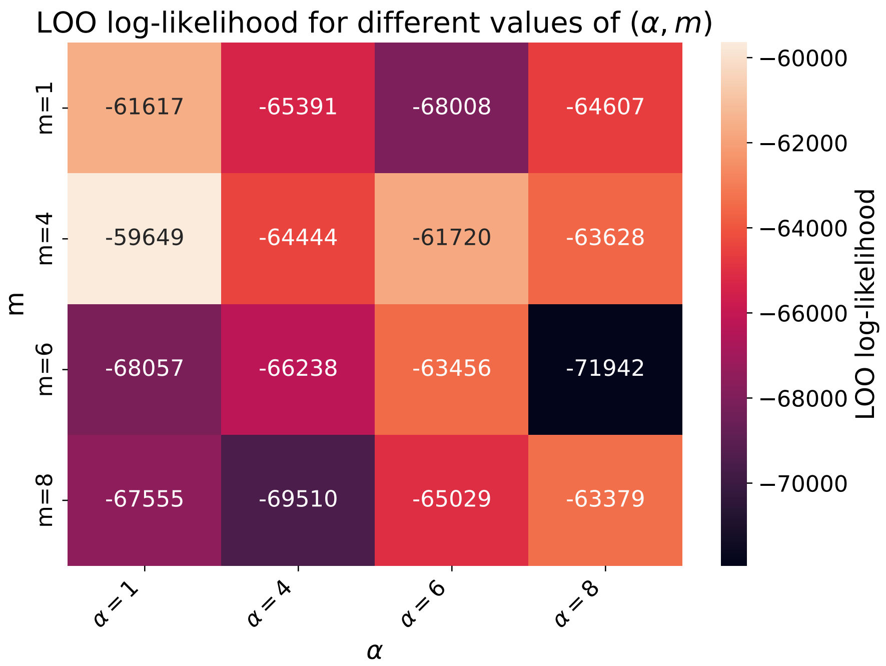


**Fig. A4**: Leave-one-out log-likelihood for the optimal number of clusters (according to posterior similarity matrix) for different values of $(\alpha, m)$.

revealed their evolutionary relationship. Genomics **103**(1), 94–106 (2014)

[4] Vehtari, A., Mononen, T., Tolvanen, V., Sivula, T., Winther, O.: Bayesian leave-one-out cross-validation approximations for gaussian latent variable models. Journal of Machine Learning Research **17**(103), 1–38 (2016)

[5] Rand, W.M.: Objective criteria for the evaluation of clustering methods. Journal of the American Statistical association **66**(336), 846–850 (1971)
